# Detection of sleep apnea from single-channel electroencephalogram (EEG) using an explainable convolutional neural network

**DOI:** 10.1101/2021.04.11.439385

**Authors:** Lachlan D Barnes, Kevin Lee, Andreas W Kempa-Liehr, Luke E Hallum

## Abstract

Sleep apnea (SA) is a common disorder involving the cessation of breathing during sleep. It can cause daytime hypersomnia, accidents, and, if allowed to progress, serious, chronic conditions. Continuous positive airway pressure is an effective SA treatment. However, long waitlists impede timely diagnosis; overnight sleep studies involve trained technicians scoring a polysomnograph, which comprises multiple physiological signals including multi-channel electroencephalography (EEG). Therefore, it is important to develop simplified and automated approaches to detect SA. We have developed an explainable convolutional neural network (CNN) to detect SA from single-channel EEG recordings which generalizes across subjects. The network architecture consisted of three convolutional layers. We tuned hyperparameters using the Hyperband algorithm, optimized parameters using Adam, and quantified network performance with subjectwise 10-fold cross-validation. Our CNN performed with an accuracy of 76.7% and a Matthews correlation coefficient (MCC) of 0.54. This performance was reliably above the conservative baselines of 50% (accuracy) and 0.0 (MCC). To explain the mechanisms of our trained network, we used critical-band masking (CBM): after training, we added bandlimited noise to test recordings; we parametrically varied the noise band center frequency and noise intensity, quantifying the deleterious effect on performance. We reconciled the effects of CBM with lesioning, wherein we zeroed the trained network’s 1st-layer filter kernels in turn, quantifying the deleterious effect on performance. These analyses indicated that the network learned frequency-band information consistent with known SA biomarkers, specifically, delta and beta band activity. Our results indicate single-channel EEG may have clinical potential for SA diagnosis.

## Introduction

Sleep apnea (SA) is a progressive disease which involves repeated episodes of apnea and/or hypopnea during sleep. It afflicts between 2% and 7% of the general population [1,2] and causes fragmented sleep. Sufferers often experience daytime hypersomnia, cognitive dysfunction that accompanies sleepiness, and an increased risk of workplace and motor vehicle accidents [3]. If allowed to progress, SA is associated with a range of serious, chronic conditions, including cardiovascular and cerebrovascular disease, and diabetes [1,2,4,5]. Continuous positive airway pressure (CPAP) -- the gold standard SA treatment -- is highly effective [4]. However, there are barriers to SA detection, diagnosis, and treatment. The primary clinical tool for SA detection and diagnosis is overnight polysomnography (PSG), which involves sleep studies and manual scoring of recorded physiological signals by trained healthcare professionals [6]. PSG is instrumentation-intensive -- it typically involves monitoring nasal or oral airflow; thoracic and/or abdominal movement; snoring; oxygen saturation; multi-channel electroencephalogram (EEG); electrooculogram (EOG); electrocardiogram (ECG); and, electromyogram (EMG) -- and therefore, may interfere with standard patterns of sleep [7]. Overnight PSG is typically followed by manually titrated CPAP therapy [1]. Demand for overnight PSG exceeds supply; wait times for those requiring screening and diagnosis can range from 2 to 60 months [8], and it is estimated that a large proportion of SA sufferers remain undiagnosed [9].

Tools to aid detection, diagnosis, and treatment of SA are therefore a worthy pursuit, especially those involving simplified instrumentation and automation, with potential for use in the home as well as the clinic [10–12]. Automated systems making use of explainable machine learning [13] could, potentially, be used to expedite and augment diagnostic and treatment decisions; in general, explainable systems are those wherein mechanisms of detection and/or classification are made available to clinicians.

The traditional approach to SA detection from EEG involves computation of features (e.g., energy and energy variance [14]) within predefined frequency bands [14,15]. Features are then concatenated to form a high-dimensional feature vector for use in classification. Convolutional neural networks (CNNs) are a form of artificial neural network, loosely inspired by hierarchical, computational models of visual processing in the cerebral cortex (review by LeCun et al. [16]). CNNs use convolution as a form of shift-invariant feature extraction, and learn, through training, to extract salient features from time series signals (or images) that are useful for classification. Recently, CNNs have demonstrated proficiency for the classification of images [17] and signals across a range of domains, including multi-channel EEG [18]. In contrast to traditional approaches, the CNN we develop here requires no postulation of features and frequency bands at the outset, meaning that features not traditionally associated with SA could be learned during the training procedure.

We hypothesized, first, that there is information in single-channel EEG enabling reliable subjectwise detection of SA by a CNN; by “subjectwise”, we mean a CNN trained using data collected from a cohort of subjects, 1 through N, should generalize to detect SA in a previously unseen subject, N+1. Second, we hypothesized that knowledge of sleep stage (e.g., rapid-eye-movement sleep) should improve this SA detection. This hypothesis is reasonable because both SA as well as sleep stage are accompanied by characteristic alterations of EEG, and SA is associated with sleep stage (a point we elaborate in the Discussion). Third, we hypothesized that the network features enabling SA detection should be consistent with known SA biomarkers. To test these hypotheses, we trained a CNN to detect SA using single-channel EEG. To explain the trained CNN’s mechanisms, we used two visualization techniques -- critical-band masking [30,31] (wherein band-limited noise was added to signals used to test the trained network) and filter lesioning [32] (wherein 1st-layer filter kernels comprising the trained network were zeroed in turn, and the effect on performance was quantified).

## Methods

### Datasets

We trained and tested our CNN using the combination of three PSG databases. The majority of data were drawn from Sleep Health Heart Study (SHHS) Visit 2, which contains overnight EEG recordings from 2,650 patients sampled at either 125 Hz or 128 Hz [19,20]. The remaining data were drawn from the University College Dublin (UCD) Sleep Apnea Database [21] consisting of recordings from 25 participants sampled at 128 Hz, and the MIT-BIH Polysomnographic Database [21,22] with usable recordings from 16 patients sampled at 250 Hz. For consistency across databases and participants, only recordings sampled from channel C4-A1 (international 10-20 system [23]) were selected for the analysis. Recordings were low-pass filtered and downsampled to 125 Hz (cutoff = 42 Hz, −6 dB at 63 Hz).

Recordings were split into labelled 30-second segments. For SHHS and UCD data, we labelled all segments that contained 10 continuous seconds of “obstructive sleep apnea”, “central sleep apnea”, “mixed apnea”, or “hypopnea” as, simply, “apnea”. All other segments were labelled “non-apnea”. For MIT-BIH data, we used the given labels. After segmentation, we standardized EEG segments through z-score normalization (i.e., zero mean and unit variance). In addition, we ensured an equal distribution of apnea and non-apnea segments by randomly undersampling the majority class of each subject. The distribution is detailed in Table 1.

**Table 1:** The distribution of annotations within the dataset before and after undersampling. Each EEG segment has an apnea annotation (i.e., “apnea” or “non-apnea”) and a sleep-stage annotation (i.e., “wake”, “REM” or “NREM”). Overall there was on average 1,144 segments per patient before undersampling and 378 segments per patient after resampling.

| Number of Annotations | Apnea | Non-Apnea | Wake | REM | NREM |
| --- | --- | --- | --- | --- | --- |
| Before | 508,168 | 2,523,647 | 1,196,060 | 354,134 | 1,481,621 |
| After | 500,469 | 500,469 | 270,491 | 210,317 | 520,130 |

### Classifier Design

We constructed a CNN to detect SA. Schirrmeister and colleagues [18], demonstrated comparable performance between shallow and deep networks when classifying multi-channel EEG recorded from experimental participants performing a motor task. In light of their results, we constructed our networks (Figure 1) using three convolutional layers; each convolutional layer involved normalization, exponential linear unit (ELU) activation, and max-pooling. We kept our network architecture simple by using a relatively low number of convolutional layers, as we anticipated that it would facilitate explainability (i.e., mechanistic analysis of the trained network’s performance, described below). We used batch normalisation (after each convolution layer) and dropout (after each pooling layer) [24] to improve network stability and safeguard against overfitting. As illustrated in Figure 1, the last convolutional layer was followed by a dense layer, and an output layer with the softmax activation function. We initialized the weights of our CNN by drawing from a truncated normal distribution with zero mean. To train the CNN, we minimized the cross-entropy loss function through backpropagation. Backpropagation was optimized using Adam [25] in the standard fashion: alpha coefficient was set at 0.9, and beta coefficient at 0.999. The learning rate was tuned alongside other hyperparameters (see Table 2).

**Figure 1 caption:**
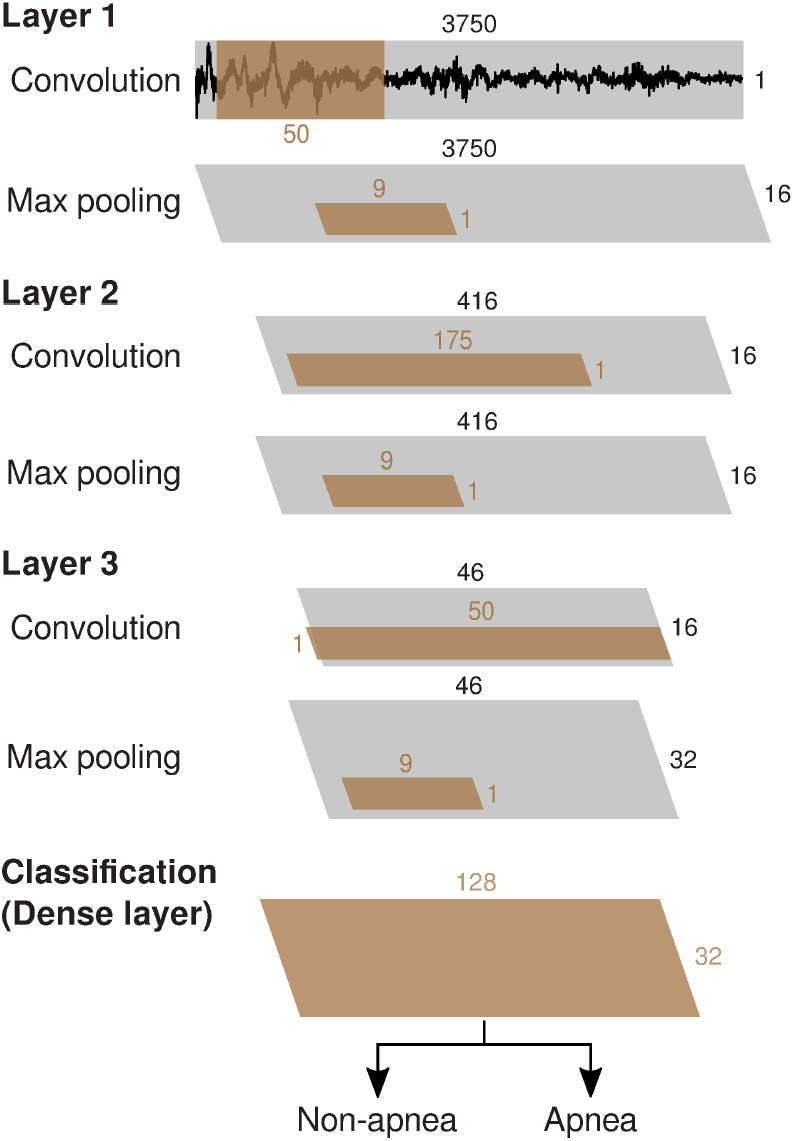
The architecture of our CNN trained to detect SA, comprising three convolutional layers. Convolutions had a stride size of one and used zero padding. Each convolutional layer was followed by batch normalisation, ELU activation, and dropout (these operations are not illustrated). The dense layer was preceded by a flattening operation, and followed by ELU activation and a dropout layer. The output layer used a softmax classifier [26]. (Symbology after Schirrmeister and colleagues [18].)

**Table 2:** Optimized hyperparameters, and hyperparameter search spaces. The rightmost column shows optimal hyperparameters evaluated with Hyperband-based tuning.

| Layer | Hyperparameter | Search space | Final |
| --- | --- | --- | --- |
| Convolutional Layer 1 | Kernel length | 25,35,50,75,125,175 | 50 |
|  | Number of filters | 8,16,32,64,128 | 16 |
| Convolutional Layer 2 | Kernel length | 25,35,50,75,125,175 | 175 |
|  | Number of filters | 8,16,32,64,128 | 16 |
| Convolutional Layer 3 | Kernel length | 25,35,50,75,125,175 | 50 |
|  | Number of filters | 8,16,32,64,128 | 32 |
| MaxPooling Layer | Window/stride size | 3,5,7,9 | 9 |
| Dense layer | Number of nodes | 16,32,64,128,256 | 128 |
| Convolutional layer dropout | Dropout rate | 0-0.6 | 0.4 |
| Dense Layer dropout | Dropout rate | 0-0.6 | 0.4 |
| Optimizer | Learning Rate | 1e-4 to 1e-1 | 0.000155 |

### Hyperparameter Tuning

We optimized the network hyperparameters (i.e., network variables set prior to training) in an automated manner. Our optimization algorithm selected hyperparameter values from a search space, and after evaluating network performance, compared the selected configuration against other combinations of hyperparameters. Suitable approaches for hyperparameter tuning are Bayesian-based [27] or Hyperband-based [28] algorithms. We tested both approaches in preliminary experiments and found that the latter outperformed the former so ultimately the Hyperband tuner was chosen. The search space and chosen hyperparameters are listed in Table 2. We adjusted Hyperband such that the algorithm was repeated five times, with one-third of hyperparameter configurations kept during its successive halving sub-operation.

### Evaluation Process

For our network, we performed subjectwise 10-fold cross-validation to assess performance. For each of these 10 folds, patients were allocated to either the training set (approx. 90%) or the testing set (approx. 10%). A randomly selected fraction of the training set (10%) was used as validation data, and was not used for training. Network training was performed for 40 epochs (i.e., 40 passes of the training data though the network). We trained and tested the CNN with Python 3.7 and Tensorflow 2.2, on NeSI (New Zealand eScience Infrastructure), a high-performance computing platform which uses Tesla P100 GPU cards (NVIDIA, Santa Clara, California, United States). Computation took 112 microseconds per input sample for training and 40 microseconds per sample for testing.

To quantify performance of our trained CNN, we used accuracy and Matthews correlation coefficient (MCC). Accuracy is a common metric used to evaluate neural network performance. However, it is susceptible to biases when data sets are unbalanced (i.e., data sets containing a preponderance of one or other class labels). Therefore, we also used MCC which is robust when data sets are unbalanced. Additionally, we developed a shuffle test to conservatively estimate performance baselines. To do so, we shuffled the labels of the training set and repeated our 10-fold subjectwise cross-validation. We reasoned that a CNN trained on these shuffled labels would be incapable of learning salient EEG features for SA detection, but could nonetheless learn the statistics of data set imbalance, and bias its behaviour accordingly. We used a Bayesian t-test, computing 95% highest density intervals (HDIs) [29], to compare our CNN’s performance to baseline.

### Critical Band Masking

To explain the mechanisms of our trained networks, we used a critical-band masking (CBM) technique that was adapted from psychoacoustics [30] and visual psychophysics [31]. Here, we added bandlimited noise to test segments (but not training segments). We used a noise bandwidth of 1.5 Hz and parametrically varied the noise frequency centred from 1.5 Hz to 60 Hz in increments of 1.5 Hz. A finite-impulse-response (FIR) band-pass filter was applied to white noise to create this bandlimited noise (length = 825, transitional bandwidth = 0.5 Hz). At each center frequency, we quantified noise intensity by computing the log of the noise root-mean-square (RMS) value and signal RMS value to find the signal-to-noise ratio (SNR). For each band center frequency, we tested the trained network using these noisy test segments, quantifying the deleterious effect of noise by observing changes in MCC scores. This CBM process was performed for every fold of our subjectwise 10-fold cross-validation (see *Evaluation Process*).

### Filter Lesioning

We adapted a “lesioning” technique from Lawhern et al. [32] to determine the relative importance of first-layer convolutional kernels to the trained network. On each fold, after having trained the network, we zeroed all coefficients for one convolutional kernel, and then tested the network. We therefore quantified the deleterious effect that zeroing (i.e., lesioning) kernels had on test performance. We did this for each first-layer convolutional kernel on each fold of our subjectwise 10-fold cross-validation. Thus, on each fold, we were able to rank 1st-layer convolutional kernels by importance; e.g., the most important kernel, when lesioned, caused the greatest reduction in network performance. To verify that this lesioning technique was effective in identifying important convolutional kernels, we computed a correlation coefficient between all pairs of kernels within and between folds; specifically, we computed Pearson’s correlation coefficient between kernels’ Fourier amplitude spectra. The correlation coefficient was generally higher between convolutional kernels deemed to be important, both within and between folds. Finally, we computed the Fourier transform, and calculated the z-scores of the most important kernels (i.e., those determined important by Filter Lesioning). To do so, we formed a null distribution of kernel transforms using all trained kernels across all folds.

## Results

Our network detected SA with an accuracy equal to 76.7% (the mean across folds of our subjectwise 10-fold cross-validation). The standard deviation (s.t.d.) of this accuracy was 0.40 percentage points across folds. Our SA network performed with an MCC score equal to 0.542 (s.t.d. = 0.009 across folds). These performance metrics were reliably above our conservative baselines (accuracy: Bayesian t-test, 0.760 < 0.767 < 0.775, 95% HDI; MCC: Bayesian t-test, 0.529 < 0.543 < 0.556, 95% HDI; *Methods*); baseline accuracy was 0.5, and baseline MCC was 0.00.

Several previous studies have found that SA is associated with sleep stage (a point we elaborate in *Discussion*). Therefore, we wondered if our SA network, to aid its performance, was representing (i.e., covertly decoding) sleep stage using sleep stage-associated features contained in these EEG recordings. To explore this idea, we decomposed our confusion matrix (Figure 2) into three submatrices, each corresponding to one or other of three sleep stages (Figure 3): wake, rapid-eye-movement (REM), and non-rapid-eye-movement (NREM). A key observation arising from inspection of these submatrices was that, for segments recorded during REM sleep, our SA network appeared to behave in a biassed fashion, predicting apnea on 95.4% of all REM trials. Our SA network also appeared to behave in a biassed fashion when detecting SA in recordings during wake; the network predicted non-apnea on 87.8% of all wake trials. These observations indicated that, potentially, our trained SA network was representing (i.e., covertly decoding) sleep stage, and, since SA is associated with sleep stage, using this representation to aid its performance in SA detection.

**Figure 2 caption:**
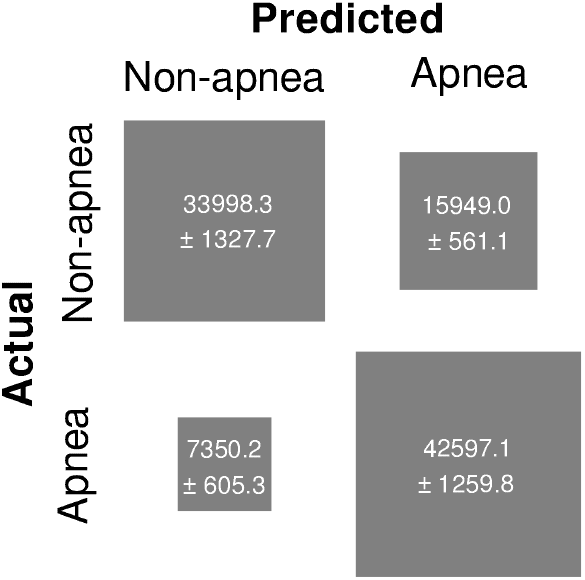
Confusion matrix, summarizing performance of our SA network. The area of each square represents the value of each matrix entry. Values are counts averaged across our subjectwise 10-fold cross-validation. The intervals (±) associated with each value show s.t.d. across folds. Overall, the network performed with accuracy = 76.7%, as indicated by the mass along the matrix’s main diagonal.

**Figure 3 caption:**
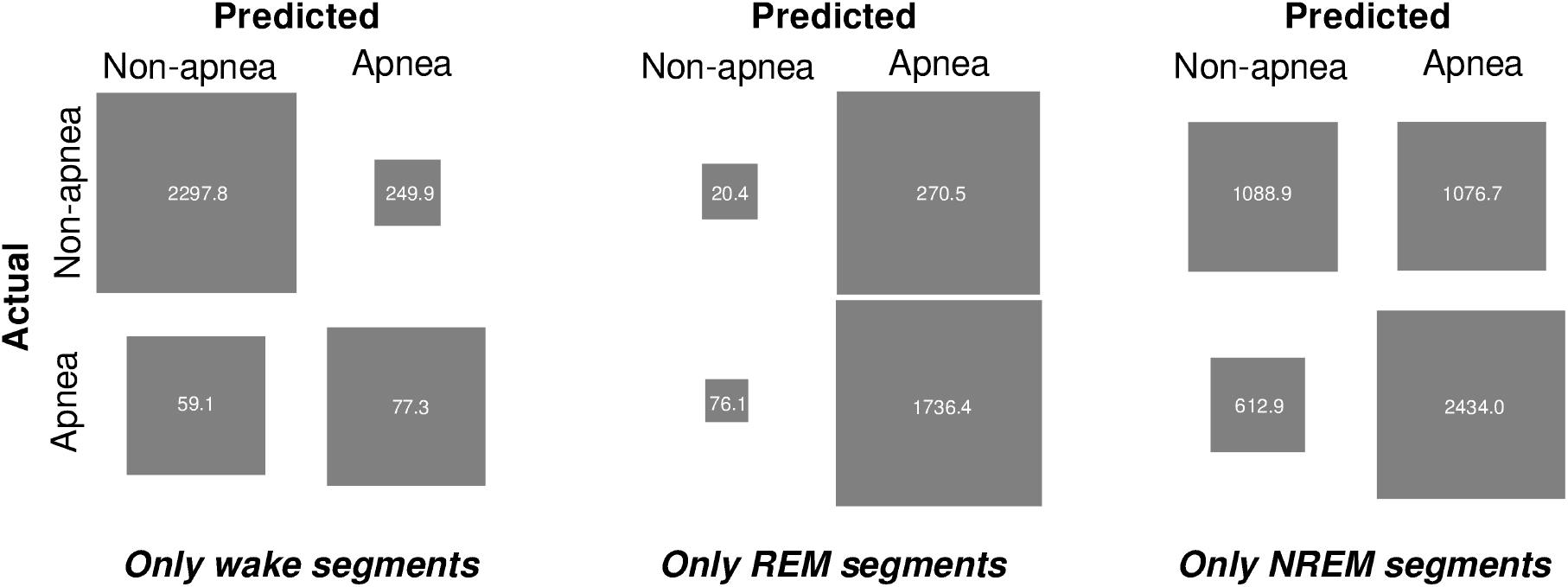
Confusion submatrices, each corresponding to one or other of three sleep stages: wake (left), REM (middle), and NREM (right). For wake and REM, our SA network appeared to behave in a biassed fashion. Graphical conventions are as in Fig. 2.

To test this hypothesis -- that our CNN was representing sleep stage -- we adapted our shuffle test (see *Evaluation Procedure*) to account for sleep stage; training data were split according to three sleep stages (wake, REM, NREM) and, for each stage, the labels “apnea” and “non-apnea” were shuffled prior to re-training the network (below, we refer to this procedure as re-training the network after a “stage-wise shuffle” of the data). We reasoned that, if our SA network was, in fact, representing sleep stage to aid its detection of SA, this stage-wise shuffle would isolate the effect of this representation on SA detection performance. Specifically, if, after we re-trained the network using stage-wise shuffled data, network performance was unaltered, then that would indicate that the representation of sleep stage was wholly responsible for SA detection. On the other hand, if, after we re-trained the network using stage-wise shuffled data, network performance fell to baseline, then that would indicate that the representation of sleep stage was not being used to aid SA detection. After re-training the network on stage-wise shuffled data, the network performed with an accuracy equal to 70.9% (the mean across folds of our subjectwise 10-fold cross-validation; s.t.d. = 0.72 percentage points across folds), and a MCC score equal to 0.466 (s.t.d. = 0.013 across folds). These performance metrics were reliably below those of the original SA network (accuracy difference to baseline: −0.064 < −0.057 < −0.050 HDI; MCC: Bayesian t-test, 0.063 < 0.076 < 0.088, 95% HDI; Methods). Therefore, our SA network appeared to learn a representation of sleep stage to aid its detection of SA, but the network’s learning of this representation only partly accounted for its ability to detect SA.

To explore further this idea -- that our SA network was representing sleep stages -- we developed a second CNN. This second CNN was nearly identical to our original SA network (the one exception being that the second CNN had five nodes in the final dense layer as opposed to two); we trained this network to decode sleep stages (wake, REM, and NREM). This sleep-stage network performed with an accuracy equal to 85.3% (mean across folds of our subjectwise 10-fold cross-validation; s.t.d. = 0.57 percentage points across folds), and a MCC score equal to 0.766 (s.t.d. = 0.009 across folds). These performance metrics were reliably above our conservative baselines (accuracy: Bayesian t-test, 0.847 < 0.853 < 0.859, 95% HDI; MCC: Bayesian t-test, 0.758 < 0.765 < 0.773 95% HDI; Methods); baseline accuracy for our sleep stage network was 52.0%, and baseline MCC was 0.00. Therefore, a network with architecture nearly identical to that of our SA network can be trained, explicitly, to decode sleep stage. This adds further support to our idea that our SA network learnt to represent (i.e., covertly decode) sleep stage, and it used this representation to aid in the detection of SA.

We wondered what EEG features were used by our SA network in performing SA detection. Therefore, we used critical-band masking (CBM), adding bandlimited noise to the signals used to test our SA network (*Methods*). The effects of CBM were graded; for the addition of low-intensity noise (SNR = 20; *Methods*), masking had little effect on network performance, regardless of the noise band’s center frequency. When we increased the intensity of noise, network performance deteriorated (i.e., MCC decreased). Deterioration was pronounced for noise in some frequency bands but not others. Overall, the effects of CBM were primarily limited to three regions (Figure 4a): frequencies less than 4 Hz (the delta band); 30 to 45 Hz (the gamma band); and frequencies running from approximately 10 to 20 Hz, encompassing alpha (8 to 13 Hz), sleep spindles (11 to 16 Hz), and the lower end of the beta band (14 to 30 Hz). Specifically, when high-intensity (SNR = 0) noise was added to the delta band, MCC for SA detection was reduced from approximately 0.54 to 0.17. When high-intensity (SNR = 0) noise was added to the band associated with sleep spindles (11 to 16 Hz), MCC for SA detection was reduced from approximately 0.54 to 0.22. When high-intensity (SNR = 0) noise was added to the band running from 15 to 18 Hz, MCC for SA detection was reduced from approximately 0.54 to 0.17.

**Figure 4 caption:**
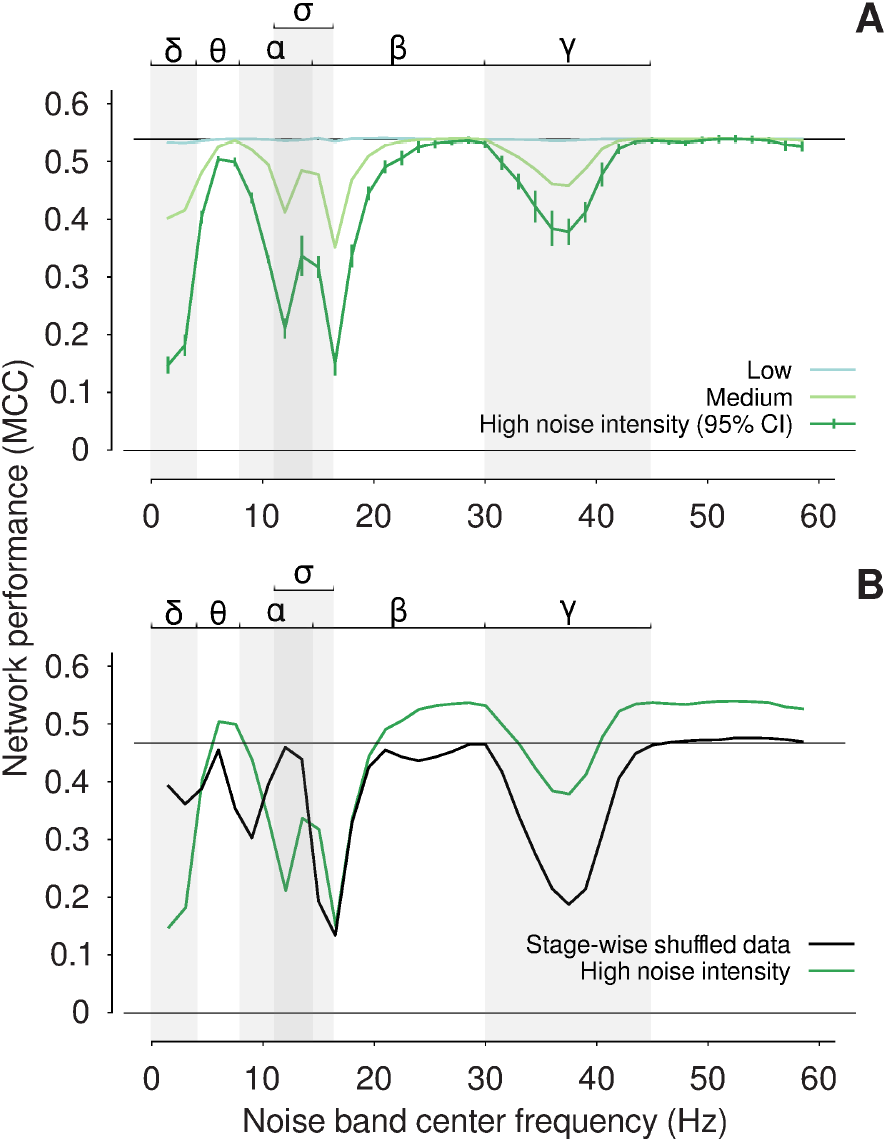
(a) Effect of critical-band masking on our SA network’s performance. We used high-, medium-, and low-intensity noise: SNR = 5, 10, and 20, respectively (*Methods*). Overall, high-intensity noise decreased performance more than low-intensity noise. The deleterious effect of noise was pronounced in some frequency bands but not others. E.g., adding bandlimited noise to test signals in the delta band (< 4 Hz) caused MCC to decrease from 0.54 to 0.17. The lower, horizontal solid line (performance = 0.0 MCC) indicates the performance baseline (*Methods*), and the upper, horizontal solid line indicates network performance in the absence of noise (performance = 0.542 MCC). The error bars (shown only for high-intensity noise) are 95% confidence intervals computed across folds of our subjectwise 10-fold cross-validation. The Greek letters (top) mark traditional frequency bands; sigma marks the band associated with sleep spindles. (b) Effect of critical-band masking on our re-trained SA network; we re-trained the network after data were stage-wise shuffled. Adding bandlimited noise to test signals in the delta and alpha (8 to 13 Hz) bands, here, had little effect on network performance. Other graphical conventions are as in (a).

For comparison, we also applied CBM to the network that we re-trained using stage-wise shuffled data (see above). We reasoned that the different effects of CBM on those two networks (our original SA network, and the network re-trained on stage-wise shuffled data) would help determine which frequency bands were important to SA detection per se, which were important to sleep stage decoding (which appears to play a role in SA detection), and which were important to both. A key outcome of this experiment involved the delta band (Figure 4b); while noise in the delta band caused deterioration in the performance of our original SA network, it had relatively little effect on the network re-trained on stage-wise shuffled data. This difference indicated that, while other frequency bands may have contributed to the network’s representation of sleep stage (e.g., sleep spindles, 11 to 16 Hz), delta band activity was specifically important to SA detection. This result is consistent with several studies that have observed an association between SA events during sleep and delta-band activity [18, 19] (a point elaborated in *Discussion*).

Critical band masking indicated that specific frequency bands were important for SA detection, especially the delta band. We therefore wondered whether filters in the first convolutional layer of our trained SA network responded selectively to EEG signals in these bands. To answer this question, we lesioned first-layer filters comprising the trained SA network (*Methods*). We found that, when lesioned, some filters substantially reduced the performance of our SA network (i.e., these filters appeared important to the network). The lesioning of other filters, by comparison, caused negligible deleterious effects (i.e., apparently unimportant filters). We sorted 1st-layer filters based on importance. These sorted filters were then used to generate an “importance matrix” (Figure 5). In light of that importance matrix we wondered, if important filters have similar characteristics? To answer this question, we computed each 1st-layer filter’s amplitude spectrum using the Fourier transform, and we averaged spectra within importance and across folds (i.e., after Fourier transformation, we averaged amplitude spectra within rows, across columns, of Figure 5). When we compared the amplitude spectra of relatively important filters to the ensemble (i.e., all filters comprising Figure 5), we found the following (Figure 6): important 1st-layer filters tended to attenuate the delta band, and amplify the beta and gamma bands. This pattern is consistent with the result of our critical-band masking.

**Figure 5 caption:**
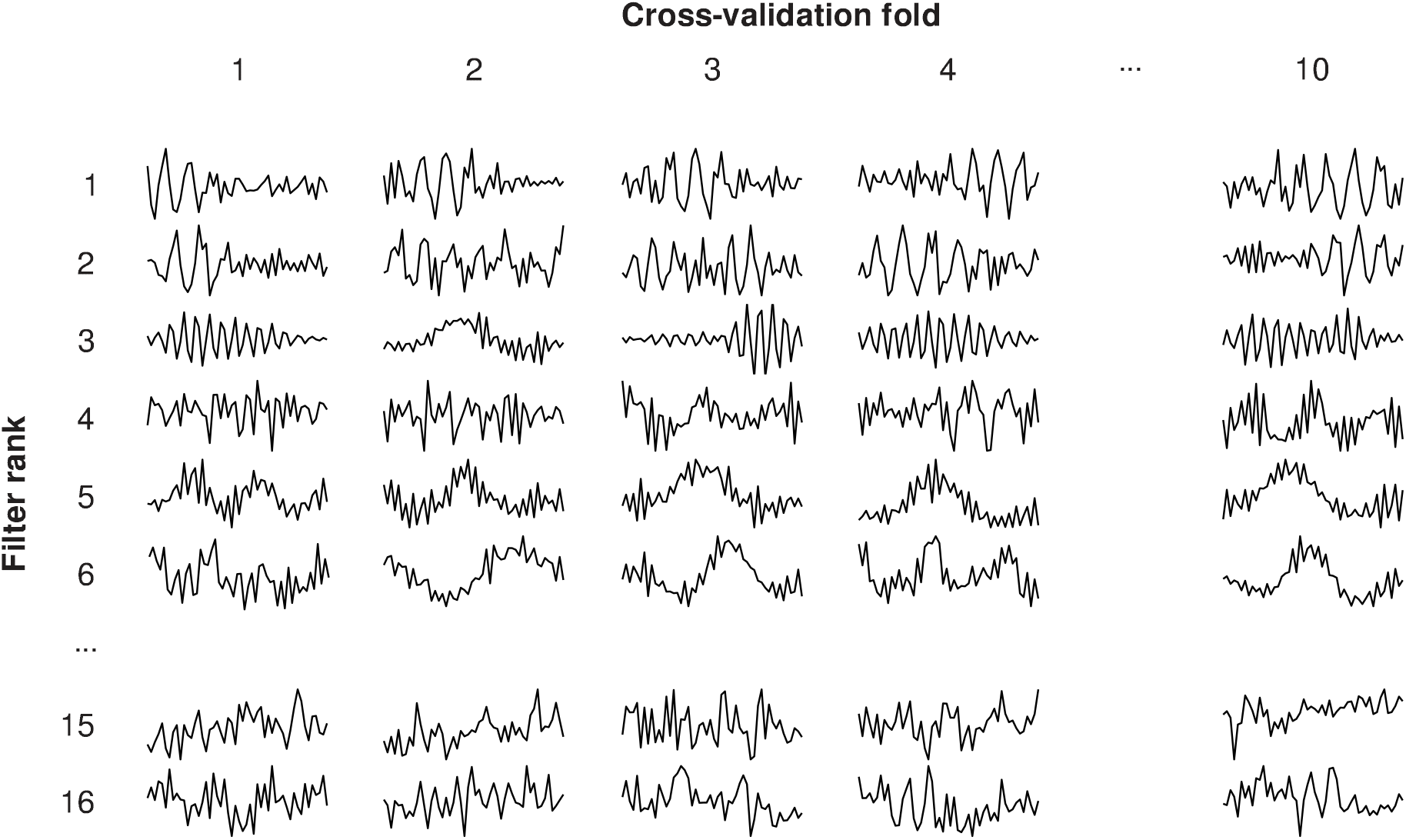
Importance matrix, showing 1st-layer filters comprising our trained SA network. Matrix columns correspond to folds from our subjectwise 10-fold cross-validation; rows correspond to importance (the most important filter on each fold is shown in row 1). To illustrate by example, on the first fold of cross-validation, the filter kernel illustrated at column 1, row 1 (top-left), was determined to be the most important; lesioning this filter reduced the trained SA network’s performance from MCC = 0.54 to 0.25. Lesioning other filters on this fold (i.e., the other filters in column 1) also reduced performance, but in all of those cases the reduction of performance was less than 0.29 (i.e., the difference between 0.54 and 0.25).

**Figure 6 caption:**
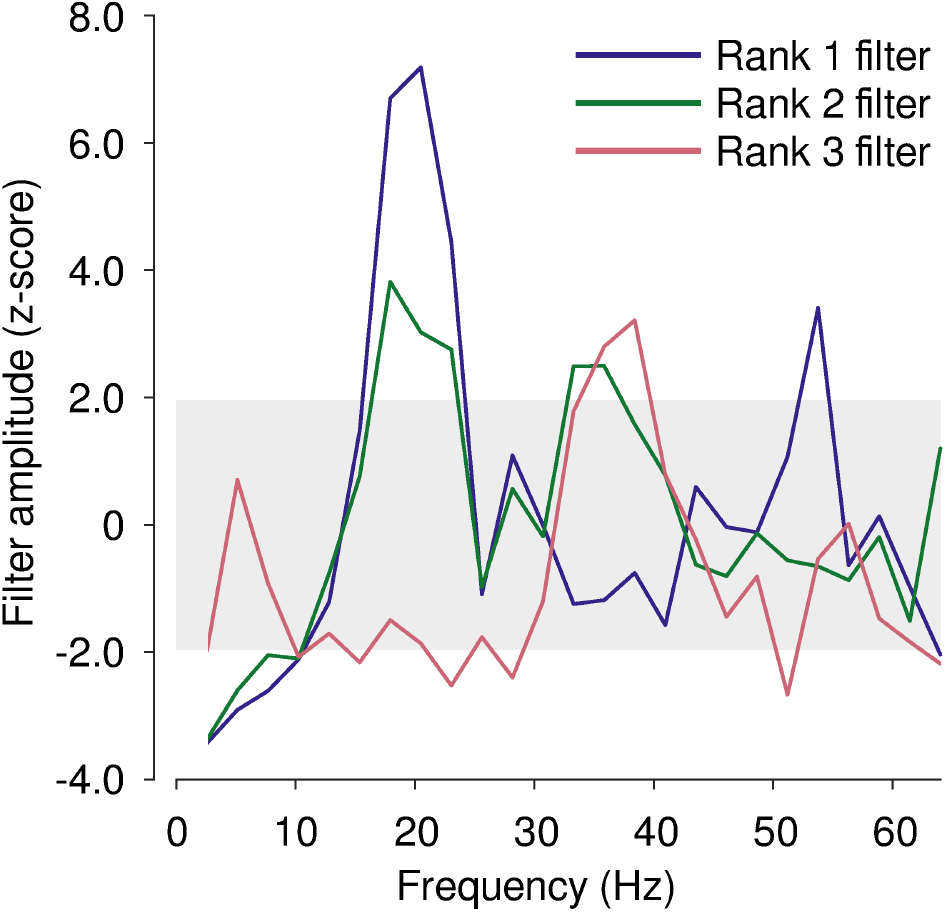
Amplitude spectra of 1st-layer filters important to the SA network’s performance. We show spectra for the 1st-, 2nd-, and 3rd-most important filters (“Rank 1, 2, and 3”, respectively). Important filters appeared to attenuate the delta band, and amplify the beta and gamma bands. The shaded rectangle marks a 95%-confidence interval (i.e., −1.96 < z-score < 1.96), wherein the spectral amplitude of rank 1, 2, and 3 filters is not appreciably different from that of all other 1st-layer filters comprising the ensemble.

## Discussion

We have developed a CNN that accurately detects SA from single-channel EEG (accuracy = 76.7% on a baseline = 50.0%). To explain the mechanisms of our CNN, we used two techniques -- critical-band masking (wherein band-limited noise was added to signals used to test the trained network) and filter lesioning (wherein 1st-layer filter kernels were zeroed in turn). Our results indicate all three of our hypotheses, which we outlined in the *Introduction*, were confirmed: there is information in single-channel EEG enabling reliable subjectwise detection of SA by an explainable CNN; knowledge of sleep stage appeared to improve SA detection; and, our CNN used information contained in single-channel EEG that is consistent with known SA biomarkers, specifically the delta and beta bands.

Previous work has demonstrated that SA is accompanied by characteristic alterations of EEG [33,34]. Likewise, the sleep stages -- wake, REM, and the three NREM stages (N1, N2, N3) -- all also are accompanied by systematic changes in EEG [35,36]. Azim and colleagues [34] studied the normalized Welch power spectral density of electrode C4-A1 from the UCD database [21]. They found that during SA events, power in the beta band (i.e., frequencies between 14 and 30 Hz) decreased (compared to pre-apnea events), before rising again after SA event termination. During different stages of sleep, the EEG features associated with SA can change. For example, apneas occurring during NREM sleep are associated with a gradual increase in delta-band activity (i.e., at frequencies < 4 Hz), followed by a decrease within that band concomitant with patient arousal and/or wakefulness [33,34]. In contrast, apneas occurring during REM sleep are associated with transient increases in delta-band activity [33,34]. REM SA events are also generally associated with small increases to beta band activities. Physiological differences between sleep stages can also impact SA. REM sleep causes the relaxation of muscle tone which is conducive to SA events [37]. Therefore, EEG -- a signal which contains characteristics for both SA and sleep stages -- should in theory be useful for detecting SA.

Our use of critical-band masking and filter lesioning, taken together, indicated that our SA network learned to rely on known SA, as well as sleep stage, biomarkers. Critical-band masking indicated that delta-band activity was important to the detection of SA; that beta- and gamma-band activity was important to the decoding of sleep stage; and that alpha-band activity may have played a role in both SA detection and sleep stage decoding. We made these inferences by adding noise, first, to signals used to test our trained SA network, and, second, to signals used to test the network after re-training with stage-wise shuffled data. Our masking results were in broad agreement with our lesioning results. The trained SA network’s most important 1st-layer filters selectively attenuated delta-band activity, and selectively amplified activity between 14 and 30 Hz (i.e., the beta band). Taken together, our findings are broadly consistent with existing work; it has been previously shown using spectral analysis of multi-channel EEG recorded from SA sufferers that delta-band activity is associated with SA events [33,34]. Furthermore, REM and NREM sleep are associated with the reduction of power in beta- and gamma bands [38].

To our knowledge, Jiang and colleagues [39] is the only other group to develop a CNN for classification of SA using single-channel EEG. We have extended their work in several ways, albeit comparison between their results and ours, for reasons outlined below, is not straightforward. First, Jiang et al. used a small dataset (the MIT-BIH Polysomnographic Database [22]), comprising recordings from only 16 participants. By contrast, our study integrated recordings from three databases, comprising over 2600 participants. Second, Jiang et al. appear to have used an unbalanced data set to train and test their network (specifically, their dataset appears to have overrepresented apnea, as opposed to non-apnea, EEG recordings); in general, the use of unbalanced data sets may bias estimates of a classifier’s performance. By contrast, we were careful to balance data before training our CNN, and we developed a shuffle test to ensure that our estimates of baseline performance were conservative. Third, Jiang et al. performed pooled (not subjectwise) cross-validation. By “pooled”, we mean that on each fold of their cross-validation procedure, recordings from all subjects were contained in both training and testing sets. Because they pooled data, it is unclear whether their results can generalize in a clinical setting (i.e., it would be clinically useful if a CNN, trained using data recorded from a normative cohort of subjects 1 through N, could be used to classify SA in recordings from a previously unseen subject N+1). By contrast, we used subjectwise cross-validation, which indicates that our results will generalize to unseen participants, and therefore may have clinical potential for diagnosis. Fourth, the network of Jiang et al. was, by contrast to ours, rather complex, and therefore ill-suited for explanation; the network employed a hybrid architecture with 4 parallel, computational branches: two shallow branches (each of which comprised three convolutional layers), and two deep branches (comprising 6 and 9 convolutional layers, respectively). By contrast, our network was simple, and this simplicity enabled an explanation of its function. It is therefore our expectation that our explainable CNN has clinical potential for the detection, diagnosis, and treatment of SA.

## Acknowledgements

We thank Hazel Glasgow and Lucia Quirke for their valuable contributions at an early stage of this research. We used high-performance computing facilities provided by New Zealand eScience Infrastructure (https://www.nesi.org.nz), which is jointly funded by collaborator institutions and the New Zealand Ministry of Business, Innovation & Employment.

